# Influence of Electronic Polarization on the Binding of Anions to a Chloride-Pumping Rhodopsin

**DOI:** 10.1101/2022.12.31.522380

**Authors:** Linda X. Phan, Victor Cruces Chamorro, Hector Martinez-Seara, Jason Crain, Mark S.P. Sansom, Stephen J. Tucker

## Abstract

The functional properties of some biological ion channels and membrane transport proteins are proposed to exploit anion-hydrophobic interactions. Here, we investigate a chloride-pumping rhodopsin (ClR) as an example of a membrane protein known to contain a defined anion binding site composed predominantly of hydrophobic residues. Using molecular dynamics simulations, we explore Cl^−^ binding to this hydrophobic site and compare the dynamics arising when electronic polarization is neglected (CHARMM36 (c36) fixed-charge force field), included implicitly (via the prosECCo force field), or included explicitly (through the polarizable force field, AMOEBA). Free energy landscapes of Cl^−^ moving out of the binding site and into bulk solution demonstrate that the inclusion of polarization results in stronger ion binding and a second metastable binding site in ClR. Simulations focused on this hydrophobic binding site also indicate longer binding durations and closer ion proximity when polarization is included. Furthermore, simulations reveal that Cl^−^ within this binding site interacts with an adjacent loop to facilitate rebinding events that are not observed when polarization is neglected. These results demonstrate how the inclusion of polarization can influence the behavior of anions within protein binding sites and thereby reveal novel mechanisms.

**Statement of Significance:** Molecular simulations based on classical (Newtonian) mechanics represent the most common method of visualizing the behavior of water and ions within channels and nanopores.
Although computationally efficient, many of the approximations required mean that these simulations often do not fully capture the complex and dynamic interactions involved. Here, we use the prosECCo force field that offers an improved electronic description whilst maintaining computational efficiency. We show that using this method to include the effects of polarization greatly influences the binding dynamics of anions to a protein binding site and yields results similar to more accurate but computationally demanding methods.

## Introduction

Halide ions (F^−^, Cl^−^, Br^−^, I^−^) are ubiquitous in biological systems (1). Amongst these, Cl^−^ is the most abundant anion and is responsible for a range of physiological processes from cell sensing and signaling to regulation of membrane potential (2). These functions are performed through a diverse range of ion channels and membrane transport proteins that preferentially bind anions, in particular Cl^−^, to facilitate their movement across the membrane (3)(4). Dysfunctional Cl- channels and transporters are known to result in a variety of disease states (channelopathies) (5) and many represent attractive therapeutic targets. However, several aspects of how anions interact with these proteins remain poorly understood.

Many studies suggest that Cl^−^ can form favorable interactions with a variety of hydrophobic interfaces, from simple aqueous/air interfaces (6) to more complex interfaces with proteins (7). In these circumstances, Cl^−^ exhibits preferential adsorption to the interfacial layer where an electrolyte solution and hydrophobic medium meet. At such an interface, Cl^−^ becomes polarized due to the anisotropy of the interface, thereby inducing a dipole not present in bulk solution. Interactions between the dipole and surrounding water molecules then compensate for the partial reduction in hydration in the interfacial layer, allowing Cl^−^ to come into direct contact with the hydrophobic interface (8). Partial removal of water from the first hydration shell of Cl^−^ is considered energetically favorable (7, 9); this phenomenon not only applies to Cl^−^ but also to ions across the Hofmeister series, i.e., F^−^ < Cl^−^ < Br^−^ < I^−^. The smaller, less polarizable F^−^ generally retains its hydration shell and is unlikely to be found at aqueous/hydrophobic interfaces whereas larger, more polarizable anions such as Br^−^ and I^−^ are more easily dehydrated and localize at the interface (9, 10). This is supported by experimental studies which indicate that the larger the halide ion, the larger the magnitude of surface adsorption to hydrophobic surfaces and hence change in water structure (11, 12).

A rapidly increasing number of high-resolution structures of channels and transporters now exist, many of which exhibit preferential interactions with anions. These include the CFTR Cl^−^ channel that is defective in Cystic Fibrosis (13), and many anion transporters (14, 15) including the NTQ chloride-pumping rhodopsin (ClR) (16). However, despite the availability of such structures, the dynamics of their interactions with anions remains under-explored in part due to the computational challenges faced in the molecular modelling of polarizable components.

The majority of molecular dynamics (MD) simulations utilize classical pairwise additive force fields with fixed point charges (17). Despite their general applicability and computational efficiency, such force fields fail to accurately capture the effects of induced polarization. Consequently, this becomes problematic for the description of systems which depend upon the behavior of anions.

The role of hydrophobic contacts with anions has been investigated in several recent simulation studies using a range of force fields (7, 9, 18). In particular, ongoing work on force field design has yielded hybrid methods that possess the computational efficiency of standard non-polarizable force fields yet contain parameterizations that better represent electronic responses within a mean field framework. One such approach is the Polarization Reintroduced by Optimal Scaling of Electronic Continuum Correction Origin (prosECCo) method (19).

prosECCo is based on the initial Electronic Continuum Correction (ECC) method which accounts for polarizability implicitly through re-scaling highly charged groups by a factor of 1/*ε_el_*^½^, where *ε_el_* represents the electronic component of the dielectric constant and is estimated as the high-frequency dielectric constant (*ε_el_* = 1.78 for water and *ε_el_* = 2 for proteins) (20–22). Furthermore, the Lennard-Jones (L-J) parameters require slight adjustments as charge scaling affects the ion-water interactions, therefore a decrease of 5% – 10% in radius is recommended to recover the correct hydration structure (21). The prosECCo force field is based on CHARMM36, and when required, incorporates these concepts systematically for a wide range of ions, proteins, lipids and sugars (19).

Another method to account implicitly for polarization in a non-polarizable force field is the empirical non-bonded fix (NBFIX) corrections applied to the CHARMM force fields that yields a more accurate model for interacting charges (21, 23), and have been shown to influence the dynamics of a Cl^−^-specific transporter, CLC-ec1 (18).

In this study, we chose to examine a microbial Cl^−^-pumping rhodopsin (ClR) (**Fig. 1A**) (PDB: 5G28, 1.6 Å resolution) (16). ClR functions as a light-driven inward Cl^−^ pump. Upon photoactivation, retinal isomerization induces a conformational change that facilitates Cl^−^ transport through the protein (24). ClR contains two Cl^−^ binding sites: the first (Cl^−^#1) is located near the retinal whilst the second (Cl^−^#2) is on a cytoplasmic loop (**Fig. 1A**) where its role in the Cl^−^ transfer pathway is to facilitate Cl^−^ release into the cytosol (16). It is this binding site that is of particular interest because Cl^−^ interacts primarily with the hydrophobic moieties within this site (**Fig. 1C**).

**Figure 1:**
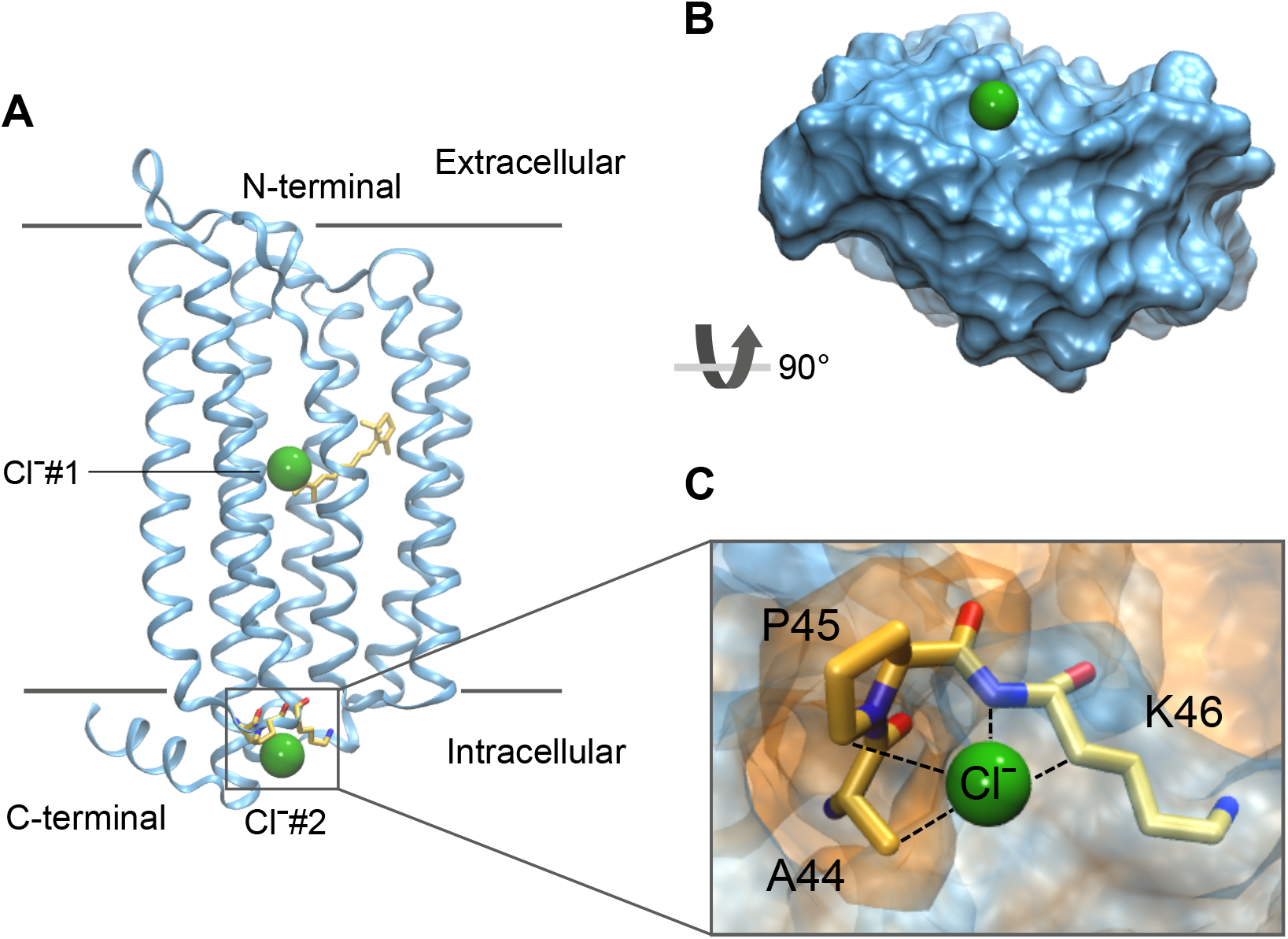
**A** Crystal structure of Cl^−^-pumping Rhodopsin (ClR) (PDB: 5G28). The first Cl^−^ binding site (Cl^−^#l) is located in the centre of the protein and is in contact with the all-*trans* retinal illustrated by the yellow stick representation in the center of the protein. The second site (Cl^−^#2) is located on the intracellular side (grey box) and is the focus of this study. The grey lines depict the extent of the lipid bilayer and Cl^−^ ions are represented by the green spheres. **B** A surface representation of the protein rotated by 90° to illustrate the bottom face where Cl^−^#2 site is located. The Cl^−^ ion is situated outside the protein. **C** A close-up illustration of the Cl^−^#2 binding site. Cl^−^#2 is comprised of the hydrophobic residues A44 and P45 (orange) and polar residue K46 (blue). The black dashed lines indicate the known contacts with Cl^−^ in the site.

Here, we performed MD simulations of ClR embedded in a lipid bilayer with the CHARMM36 (c36) and prosECCo force fields. Potential of mean force (PMF) calculations enabled us to examine the free-energy landscapes of a Cl^−^ moving out of the Cl^−^#2 site into bulk. We also simulated a smaller protein fragment that mimics this site to probe the dynamics of binding at the Cl^−^#2 site using the fully polarizable force field, AMOEBA, and compared the behavior of Cl^−^ using the prosECCo and c36 force fields. Finally, we explored how residues in a neighboring loop facilitates the rebinding of Cl^−^. Our results demonstrate that accounting for polarizability yields fundamentally different Cl^−^ interactions with ClR thereby highlighting how even simplified descriptions can replicate the behavior of anions in simulations.

## Methods

### Structural model and system preparation

Starting from the experimentally determined protein structure (PDB ID: 5G28) (16), all ligands including retinal, were removed. It was not necessary to parameterize and simulate retinal for the purposes of this study. Systems with the whole protein and/or reduced fragments containing only the binding site of interest were prepared. Simulations involving the whole protein were embedded in a 1-palmitoyl-2-oleoyl-*sn*-glycerol-3-phosphochline (POPC) bilayer and prepared using the CHARMM-GUI protocol (25–27). We chose not to apply position restraints to the whole protein simulations whereas position restraints with a force constant of 1000 kJ/mol/nm^2^ were applied to the C_α_ atoms for the reduced proteins simulations. Analysis of RMSDs suggested that the protein simulations without backbone position restraints deviated no more than 1.5 Å from the crystal structure (**Fig. S1**).

### Molecular Dynamics Simulations

For comparison, non-polarizable atomistic simulations were performed using CHARMM36 (c36) with associated lipid parameters where relevant (28). Simulations with the ECC-scaling used the prosECCo force field, a variant of the same CHARMM36 force field but with scaled charges and compatible ion types (https://gitlab.com/sparkly/prosecco/prosECCo75) (19). Whilst the theoretical framework of NBFIX has proven to successfully reproduce experimental observations through preventing overbinding (18), we chose prosECCo because it does not require additional pair-specific parameterizations of L-J parameters and has previously been applied to the study of halides (22, 29).

The systems were equilibrated through a staged protocol whereby structural restraints on the protein backbone and Cl^−^ were gradually weakened over a 15 ns equilibration period. Subsequently, all production runs of 55 ns were performed with an integration time step of 2 fs. All systems were solvated with ~ 0.15 M NaCl using the SPC/E water model (30). The temperature was maintained at 303.15 K with a coupling constant of 1.0 ps using the Nose-Hoover thermostat. Pressure was maintained at 1 bar using the Parrinello-Rahman barostat with a coupling constant of 5.0 ps. Semi-isotropic pressure coupling was used for systems containing a bilayer and isotropic pressure coupling for systems of proteins in solvent. The Verlet cutoff scheme (31) was applied and electrostatics were treated with the particle mesh Ewald method (32). The LINCS algorithm was used to constrain h-bonds only (33). Three independent repeats were carried out for each system. These simulations were performed in GROMACS (34) version 2020 (www.gromacs.org) and analyzed using MDAnalysis (35, 36).

### AMOEBA Force Field Simulations

Polarizable atomic multipole simulations were carried out in OpenMM 7.4.2 (www.openm.org). All components were modelled with the AMOEBA polarizable force field using the AMOEBA13 protein parameter set (37) and the AMOEBA03 water model (38). Starting configurations for these simulations were taken from the final frames after 15 ns of equilibration using c36 as described above. Simulations were then set up using a similar method to that described previously (https://github.com/Inniag/openmm-scripts-amoeba) (7). Simulations using the AMOEBA force field were performed only for the small and large fragments in solvent therefore all C_α_ atoms were placed under a harmonic restraint with force constant 1000 kJ/mol/nm^2^ to prevent the reduced structures deviating from the experimentally determined structure. Production runs lasted 55 ns and time integration was achieved using the r-RESPA method with an outer time step of 2 fs and an inner time step 0.25 fs. The temperature was maintained at 303.15 K using the Andersen thermostat and the pressure was maintained at 1 bar using the isotropic Monte Carlo barostat. Three independent repeats were performed for each system.

### Umbrella Sampling

This was performed to obtain one-dimensional potential of mean force (PMF) profiles for a Cl^−^ ion moving from the binding site into bulk solution. These simulations were carried out on the whole protein embedded in a POPC bilayer and solvated with ~ 0.15 M NaCl solution using the c36 and prosECCo force fields and the SPC/E water model. Systems were simulated with and without the use of backbone restraints. The collective variable (CV) was defined as the distance between the ion and the center of mass (COM) of the protein binding site (residues 44 to 46) in the negative direction parallel to the z-axis of the simulation box. Starting configurations for the umbrella windows were obtained from equilibrated simulations. The target ion was relocated to subsequent positions parallel to the z-axis and the lateral positions of the ion were set to that of the binding site COM. During equilibration and sampling, a harmonic biasing potential of 1000 kJ/mol/nm^2^ was applied to restrain the collective variable. Umbrella windows covered the distance from the ion binding site into the bulk water regime. This corresponded to 50 windows with a 0.5 Å distance interval between two successive windows. Following ion relocation, 10 steps of energy minimization were performed to remove any steric clashes between the target ion and surrounding atoms. A further 1 ns of isothermal-isobaric equilibration was simulated per window. Each umbrella window was simulated for 1 ns with c36 and 2.5 ns with prosECCo. Simulation details were similar to those detailed above for MD simulations. Unbiasing was performed through WHAM: the Weighted Histogram Analysis Method using the Grossfield lab implementation in version 2.0.9 (http://membrane.urmc.rochester.edu/wordpress/?page_id=126). PMF profiles were shifted so that the ion in bulk solution corresponds to 0 kJ/mol.

## Results & Discussion

### Protein Binding Site and Simulations

In selecting a system to study, we wished to explore anion-hydrophobic interactions within a biologically relevant protein structure. The second Cl^−^ binding site (Cl^−^#2) of ClR is located on the cytosolic surface (**Fig. 1B**) where it facilitates Cl release into the cytosol and has unobstructed access to the bulk cytoplasm (16). In this site, Cl^−^ is coordinated by an aliphatic hydrogen of A44, a hydrogen from the aromatic ring of P45, a hydrogen from the sidechain of K46 and is considered to be hydrogen-bonded to the backbone nitrogen of K46 (**Fig. 1C**).

In this study, we performed simulations of the whole protein structure with non-polarizable force fields (c36 and prosECCo). However due to the methodological and computational demands of implementing a fully polarizable force field (AMOEBA) on these larger simulations, we therefore designed a reduced system containing only the binding site of interest and a few surrounding residues that neither interact with, nor obstruct the Cl^−^ pathway. We truncated the structure to form a smaller reduced fragment consisting only the binding site of interest (residues 44 to 46) (**Fig. 1C**) and a few extra surrounding residues (residues 42 to 49). We refer to this model as the ‘smaller fragment’. A larger reduced protein system also included additional nearby residues of interest (residues 252 to 256) (**Fig. 5A**) and is referred to as the ‘larger fragment’. For these reduced protein systems, position restraints were applied to the C_α_ atoms to ensure the structures did not significantly deviate from the experimental coordinates.

**Figure 2:**
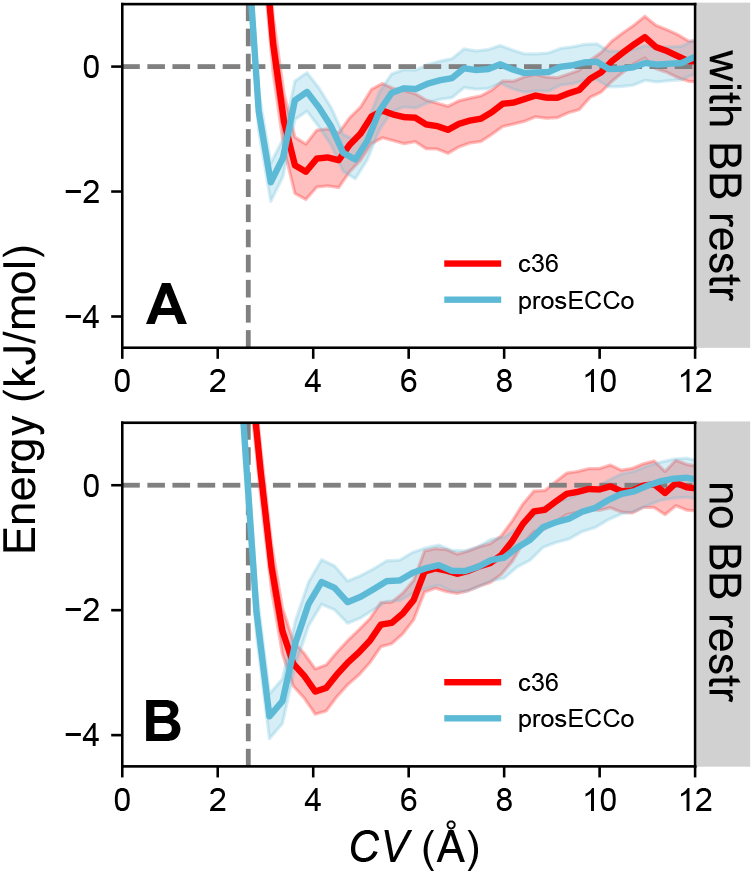
Potential of mean force profiles of a Cl^−^ ion moving away from the binding site into bulk solution. The collective variable (*CV*) is defined as a straight line parallel to the z-axis starting from the center of geometry (COG) of the binding site and into bulk solution. Free energy profiles from simulations with (**A**) and without (**B**) the application of protein backbone restraints are shown in red and blue for the CHARMM36 and prosECCo force fields respectively. Confidence bands were obtained by calculating the standard error over 200 ps and 500 ps sampling blocks during the same period for c36 and prosECCo respectively. The horizontal grey dashed line represents the free energy of a Cl^−^ ion in bulk solution. The vertical grey dashed line indicates the location of the Cl^−^ ion bound in the experimentally determined crystal structure in terms of *CV*.

**Figure 3:**
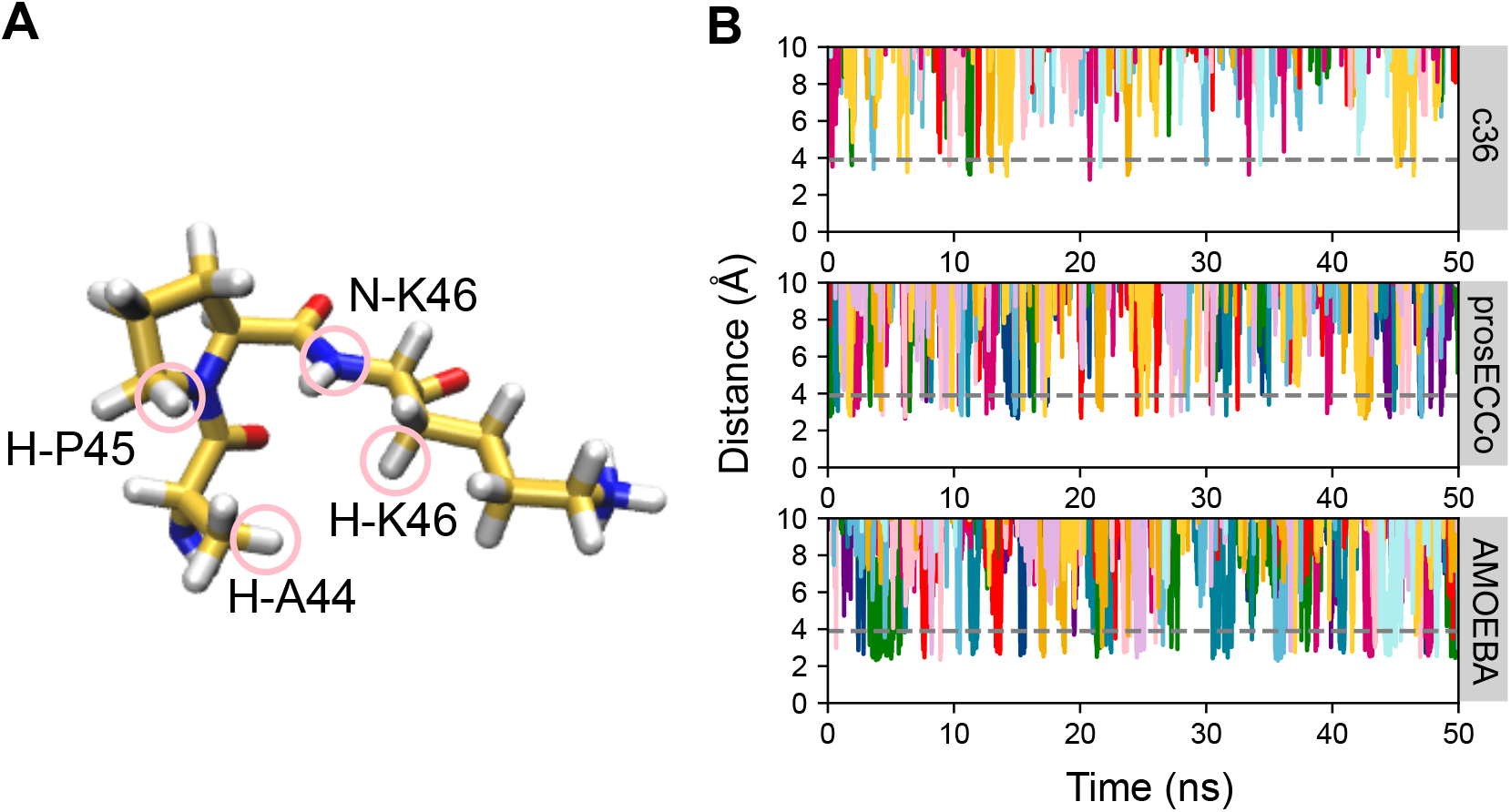
**A** Schematic diagram of hydrophobic binding site (Cl^−^#2). The atoms circled in pink correspond to the atoms used for Cl^−^ ion-distance calculations. **B** Ion-distance plots from Cl^−^ ions to an aliphatic hydrogen atom of A44 over 55 ns within simulations of the smaller protein fragment. Each color represents an individual ion trajectory, plotting only the ions that initially come into proximity of 5 Å to the binding site. The grey dashed line indicates the binding distance taken as the Cl^−^–oxygen atom of water i.e., the distance of the first hydration shell ~ 3.9 Å.

**Figure 4:**
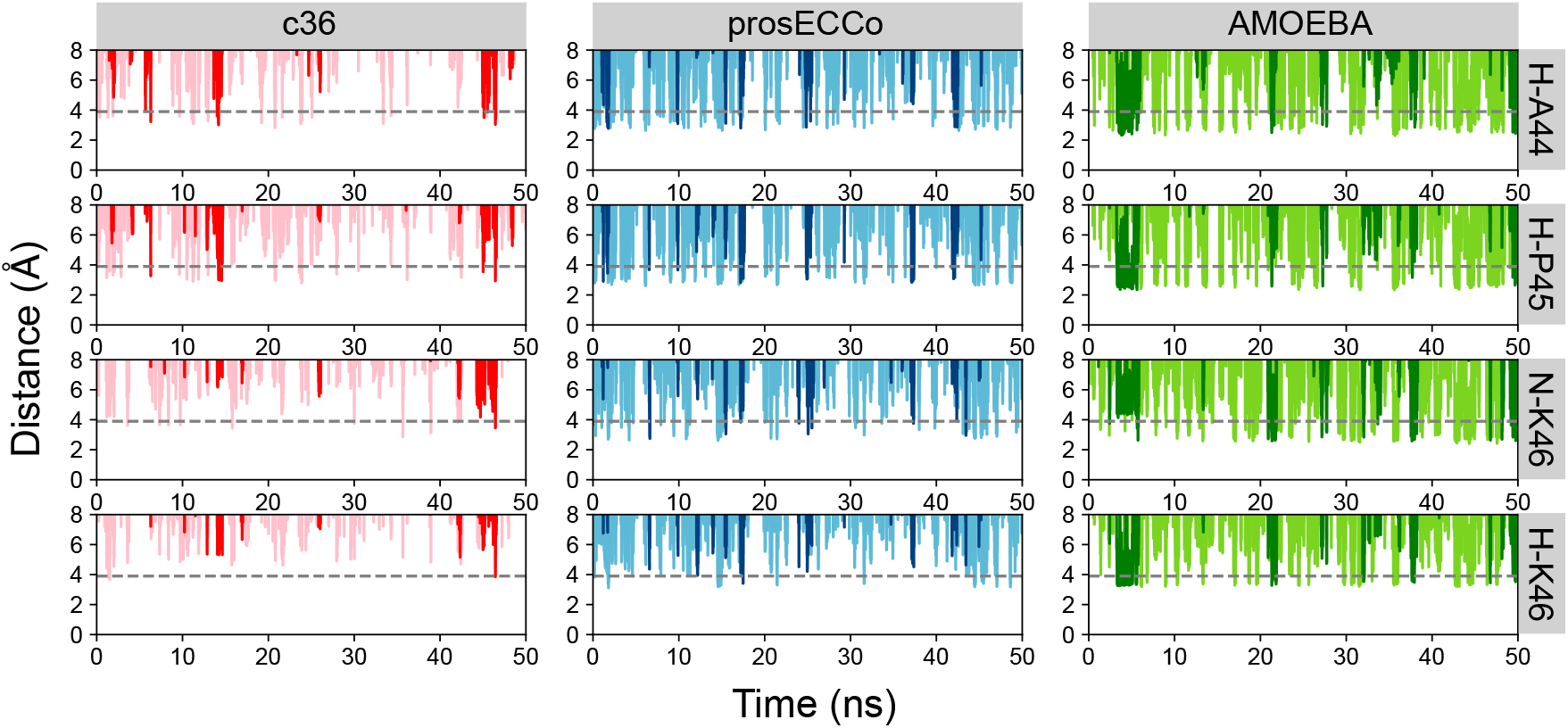
Examples of single Cl^−^ ion-distance plots (darker colors). Other Cl^−^ ion-distance plots are also shown as a function of time (lighter colors). The ion distances are measured between a Cl^−^ ion to an aliphatic hydrogen atom of A44 (H-A44), a hydrogen of the aromatic ring of P45 (H-P45), the backbone amide nitrogen of K46 (N-K46) and a hydrogen from the sidechain of K46 (H-K46) (see **Fig. 3A**). The grey dashed line indicates the binding distance taken as the Cl^−^-oxygen atom of water i.e., the distance of the first hydration shell ~ 3.9 Å.

**Figure 5:**
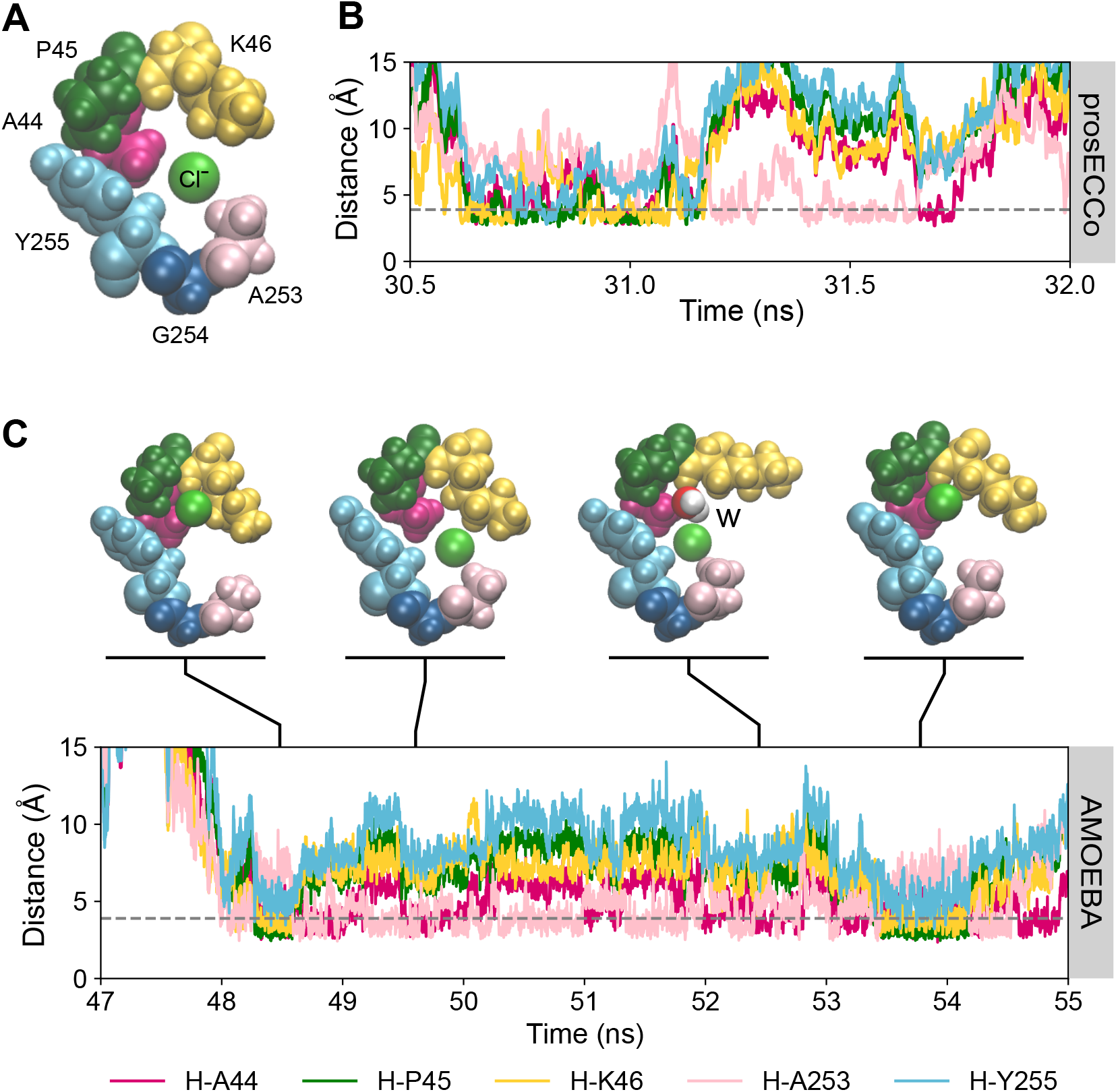
**A** Larger protein fragment containing the Cl^−^#2 binding site and residues from a nearby loop. The binding site is composed of A44 in magenta, P45 in dark green and K46 in yellow. The additional residues of interest from the nearby loop are illustrated by A253 in pink, G254 in navy and Y255 in light blue. **B** Example of an ion-distance plot as function of time for a Cl^−^ ion in the binding site simulated using the prosECCo force field. **C** An ion-distance plot as a function of time for a Cl^−^ in the binding site simulated using the AMOEBA force field. Snapshots of a Cl^−^ ion in the binding site illustrate the ion dissociation and rebinding mechanism. A water molecule, marked by ‘**w**’, can be observed to displace the Cl^−^ ion from Cl^−^#2 whilst the ion favourably interacts with nearby hydrophobic residues A253 and Y255.

### Influence of Effective Polarization on the Energetics of Cl^−^ Ion Binding

We first examined how the energetics of Cl^−^ were impacted by the inclusion of effective polarization by means of the prosECCo force field compared with standard c36. We define the collective variable (*CV*) as the z-distance from the center of geometry (COG) of the binding site. Additionally, we compared how the energetics vary when applying backbone restraints. Examination of the umbrella histograms revealed good overlap between windows indicating that the resulting potential of mean force (PMF) profiles had converged (**Fig. S2**).

The PMF profiles with backbone restraints using the prosECCo force field reveals a free energy minimum of ~ −1.9 kJ/mol at *CV* ~ 3 Å, a distance of ~ 0.6 Å further from the binding location of the Cl^−^ ion in the crystal structure (**Fig. 2A**). With c36, there is an energy well of ~ −1.7 kJ/mol situated at *CV* ~ 4 Å which corresponds to a distance of ~ 1.3 Å relative to the experimentally determined ion location. Furthermore, the prosECCo free energy profile exhibits a distinctive second local free energy minimum at *CV* ~ 5 Å with a free energy ~ −1.5 kJ/mol. This second local minimum is less pronounced when backbone restraints are not implemented (**Fig. 2B**) yet is within error of the PMF profile with backbone restraints and is not observed with c36. On closer inspection of the Cl^−^ ion at this position, this second energetic well (**Fig. 2A**) appears to be associated with the displacement of the Cl^−^ ion from the binding site by a water molecule. The ion then proceeds to form contacts with the methyl group of A253 and hydrogen atoms from the aromatic ring of Y255.

When the backbone restraints are relaxed to enable protein flexibility and unrestrained dynamics different free energy landscapes are observed. For the prosECCo force field, a free energy minimum of ~ −3.7 kJ/mol relative to bulk solution at *CV* ~ 3 Å can be observed which corresponds to a distance of ~ 0.6 Å relative to the crystal structure (**Fig. 2B**). However, this binding position is consistent with the location of the first minima of the PMF profile with backbone restraints which suggests that the shape of the binding site does not deviate much from the crystal structure when the ion is bound. A profile shift can be seen with the c36 force field which exhibits an energy well of ~ −3.3 kJ/mol relative to bulk solution at *CV* ~ 4 Å (**Fig. 2B**), ~ 1.6 Å from the crystal structure. Here, the location of the free energy minimum corresponds to a dissociated Cl^−^ ion forming favorable interactions with a nearby lipid head group, hence the crystallographic binding site goes undetected with the c36 non polarizable force field.

In both cases (with/without backbone restraints), the interactions with Cl^−^ occur more closely to the crystal structure ion interaction site when employing prosECCo than c36 which could be associated with partial ion dehydration in the site (discussed later, **Table 1**). However, we find approximately two-fold stronger binding when no backbone restraints are applied (**Fig. 2B**), an effect that occurs with both force fields.

**Table 1:**

|  | Average Hydration Number |  |  |  |  |
| --- | --- | --- | --- | --- | --- |
| Force field | H-A44 | H-P45 | N-K46 | H-K46 | Bulk |
| **c36** | 6.9 | 6.7 | 6.3 | 6.7 | 7.4 |
| **prosECCo** | 5.0 | 4.9 | 3.9 | 4.3 | 6.4 |
| **AMOEBA** | 5.2 | 5.0 | 4.3 | 5.5 | 7.3 |

ClR exhibits similar pumping activity with Cl^−^ and Br^−^, and so the anomalous signal of Br^−^ was used to locate the ion-binding sites (16). We therefore also performed PMF calculations for Br^−^ ion moving out of the binding site into bulk using prosECCo. Comparing these PMF profiles (**Fig. S3**), we observe that the larger, more polarizable Br^−^ ion binds in similar locations as the Cl^−^ ion and with similar relative free energies of binding thus supporting these experimental observations.

### Influence of Polarization on Ion Binding Dynamics

We next examined the dynamics of the ion binding using the c36, prosECCo and AMOEBA force fields applied to the smaller protein fragment. We analyzed the distances between Cl^−^ and selected atoms in the binding site that are considered to stabilize binding (**Fig. 3A**) (16). Ions within an initial radius of 5 Å of the center of geometry of the binding site were first selected and then distances to binding site atoms of selected ions were then calculated. This initial distance was appropriately chosen based on halide interactions within other protein binding sites (1). We considered Cl^−^ to be bound if it resided within the distance defined by the first hydration shell (~ 3.9 Å for all force fields). This was realized through calculating the radial distribution function (RDFs) between the Cl^−^ and surrounding oxygen atoms from water molecules in bulk solvent (*gCl^−^-O(r)*) (**Fig. S4**). In this section we focused only on distances between Cl^−^ and an aliphatic hydrogen atom from A44 (**Fig. 3A**).

For simulations with the c36 force field (**Fig. 3B**), Cl^−^ occasionally approached the binding site to interact momentarily before dissociating back into solution. These interactions were brief with an average duration of < 0.5 ns and a minimum interaction distance of ~ 3.0 Å, just within the first hydration shell distance (**Fig. 3B**, grey line). In comparison, significantly more Cl^−^ ions come into proximity of the binding site when prosECCo was used. The dynamics remain relatively transient with Cl^−^ spending ~ 0.5 ns interacting with A44 in the binding site at a minimum distance of ~ 2.9 Å. With the AMOEBA forcefield, a similar number of ions come into proximity of the binding site to interact with an even closer minimum distance of ~ 2.5 Å and longer duration of occupation ~ 2 ns.

This data suggests that polarizability, leads to stronger interactions between Cl^−^ ions and A44 in comparison to standard fixed charge models. The short interaction times observed here are expected because the functional role of this binding site is to aid ion release into the cytosol (16). Simulations and analysis involving the whole protein embedded in a POPC bilayer with the c36 and prosECCo force fields qualitatively support these conclusions (**Fig. S5**).

### Cl^−^-Hydrophobic Interactions within the Binding Site

We expanded the analysis of the small fragment simulations further to examine the interactions of Cl^−^ with the whole binding site by considering the other atoms that contribute to this site (16). Therefore, we computed the distances between the ion and hydrogen atoms from A44 (H-A44), P45 (H-P45) and K46 (H-K46) as well as the backbone amide nitrogen from K46 (N-K46) (**Fig. 3A**). This analysis was prepared for each force field and enabled us to gain a more comprehensive understanding of the mechanisms behind Cl^−^ binding to this hydrophobic site. Overall, a notable trend was found across all force fields: the Cl^−^ comes into closest proximity with the hydrophobic contacts of the binding site, namely H-A44 and H-P45 (**Fig. 4**) which are approximately equidistant. This suggests that it is more favourable for Cl^−^ to associate with these hydrophobic contacts (8). These Cl^−^ preferences were achieved through the partial loss of hydration shell; an effect that was previously observed for Cl^−^ ions in a model hydrophobic nanopore (9). As a consequence of including polarization and the resulting induced dipoles, the balance between ion-water and ion-protein interactions therefore shifts towards favouring the interactions with the hydrophobic protein contacts over water. This can be realized by calculating the average hydration numbers around Cl^−^ ion within ~ 3.9 Å of the selected binding site atoms for each given force field (**Table 1**).

For c36, the average hydration number remains similar for each hydrophobic contact. A slight reduction is observed compared to Cl^−^ in bulk solution. This retention of hydration shell correlates with the ion-distance plots (**Fig. 4**) where Cl^−^ ions do not come into as close proximity with the binding site contacts compared with prosECCo and/or AMOEBA. With the prosECCo force field, a substantial Cl^−^ ion dehydration occurs in the binding site relative to bulk solution. Based on previous studies using the ECC method (9), this effect is anticipated and is again reflected in the Cl^−^ approach distance illustrated in **Fig. 4**. Moreover, this dehydration is energetically advantageous as it promotes ion binding (**Fig. 2**). The largest dehydration is observed in simulations with the AMOEBA force field where Cl^−^ loses ~ 2 water molecules relative to bulk when in the vicinity of hydrophobic contacts. This enables Cl^−^ to exist in a more stably bound state reflected by the longer binding durations (**Fig. 4**) and increased ion occupancy (**Fig. 3B**). This is also consistent with the interactions and behaviours associated with anion hydration that have been more thoroughly explored studies by Rempe *et al*. (39, 40).

The ion-distance plots (**Fig. 4**) suggest that N-K46 is further than H-A44 and H-P45 from Cl^−^ within the site. This is a result of the tendency for the tail of K46 to enclose the anion such that N-K46 is then positioned deeper inside the concave cavity formed by the binding site as a whole (**Fig. S6**).

### Influence of Polarization on the Mechanism of Cl^−^ Binding

Based on findings from the PMF profiles discussed above (**Fig. 2**), we further explored the mechanisms that give rise to the metastable free energy minimum observed in simulations employing the prosECCo force field (second minima at ~ 5 Å, **Fig. 2A**). On closer observation of the umbrella sampling windows, the residues A253 and Y255 in a nearby loop appeared to coordinate Cl^−^ at the location of the second free energy minimum. Notably, these residues are hydrophobic. To explore the influence of these hydrophobic loop contacts, we further simulated a larger protein fragment consisting of the previous smaller fragment with an additional fragment consisting of residues from the nearby loop (residues 252 - 256). This larger fragment was then simulated using the c36, prosECCo and AMOEBA force fields. We calculated the distance between Cl^−^ and the coordinating binding site atoms as before, but also now calculating the distance to a hydrogen from the methyl group of A253 and a hydrogen of CD1 belonging to the aromatic ring of Y255 (**Fig. 5A**).

Simulations of this larger fragment with the c36 force field did not show any significant interactions with residues Y255 and A253. However, with the prosECCo force field (**Fig. 5B**) we observed instances where Cl^−^ was initially bound to its binding site then dissociated to interact with the nearby loop (e.g. A253) and then proceeded to rebind to its site once more. Due to the intriguing nature of this observation, we therefore repeated the simulations with a fully polarizable force field.

With AMOEBA, the ion also dissociated from the binding site in exchange with a water molecule (e.g., **Fig. 5C**, snapshot marked **w**). From here, the ion could then form favorable interactions with A253 and/or Y255 to remain in close proximity to the binding site thus providing the opportunity to rebind or otherwise dissociate into bulk solution. An example of such a rebinding event is illustrated in **Fig. 5C**. These findings are supported by simulations of the full protein embedded in a lipid bilayer with the c36 and prosECCo force fields where the effect was not observed with c36 but was with prosECCo (**Fig. S7**). This mechanism is therefore only observed when polarizability is considered. It is interesting to note that the ion binding/rebinding occurs on a longer timescale with AMOEBA in comparison to c36 and prosECCo suggesting that the rate of transport of Cl^−^ may be lower than that predicted by non-polarizable force fields.

Improving common fixed charged methods to better capture the effects of polarization (e.g. prosECCo) preserves computational efficiency. As demonstrated here, prosECCo can act as a proxy for identifying potentially interesting anion interactions. These can then be examined further by performing simulations using explicitly polarizable models such as AMOEBA (37), CHARMM Drude (41), etc. To some degree, prosECCo can replicate the qualitative behavior produced by more advanced force fields, however, to gain a higher level of physical accuracy, extensive simulations employing fully polarizable force fields may be required or even ab initio MD calculations.

## Conclusion

We have performed simulations of a Cl^−^-pumping rhodopsin to investigate the effects of polarizability on the binding of anions within a defined binding site. Our results demonstrate how the inclusion of electronic polarization leads to stronger anion binding events and longer binding durations. These effects result from energetically favorable interactions with a site-adjacent loop and partial ion dehydration to interact with the hydrophobic moieties of the binding site. Crucially, these binding mechanisms were only observed when polarization (effects) were incorporated into the simulations.

Future work might explore more complex biological anion binding sites by applying both prosECCo and AMOEBA force fields which will help evaluate the applicability of simplified models for complex interactions arising from polarization. Structures of future interest might include the Na^+^/I^−^ symporter (NIS) (PDB: 7UV0) that mediates active I^−^ transport and contains a highly conserved I^−^ binding site consisting of predominantly hydrophobic and aromatic residues (15). Similarly, the Cl^−^-selective human bestrophin channels (hBest1& hBest2) contain a set of highly conserved hydrophobic residues within the neck of the permeation pathway (42). These structures suggest not only an important role for hydrophobic contacts in the ion permeation pathway but potentially influential roles of anion-π interactions.

## Author Contributions

LX-P VCC and HM-S performed research and analyzed data. All authors designed research and wrote the paper. SJT, JC and MSPS obtained funding for the project.

## Acknowledgements

This work was supported by grants from the BBSRC and the EPSRC and by an EPSRC iCASE studentship award in collaboration with IBM Research to LXP. It was also supported by the Hartree National Centre for Digital Innovation - a collaboration between the Science and Technologies Facilities Council and IBM.

## Supplemental Information

**Figure S1:**
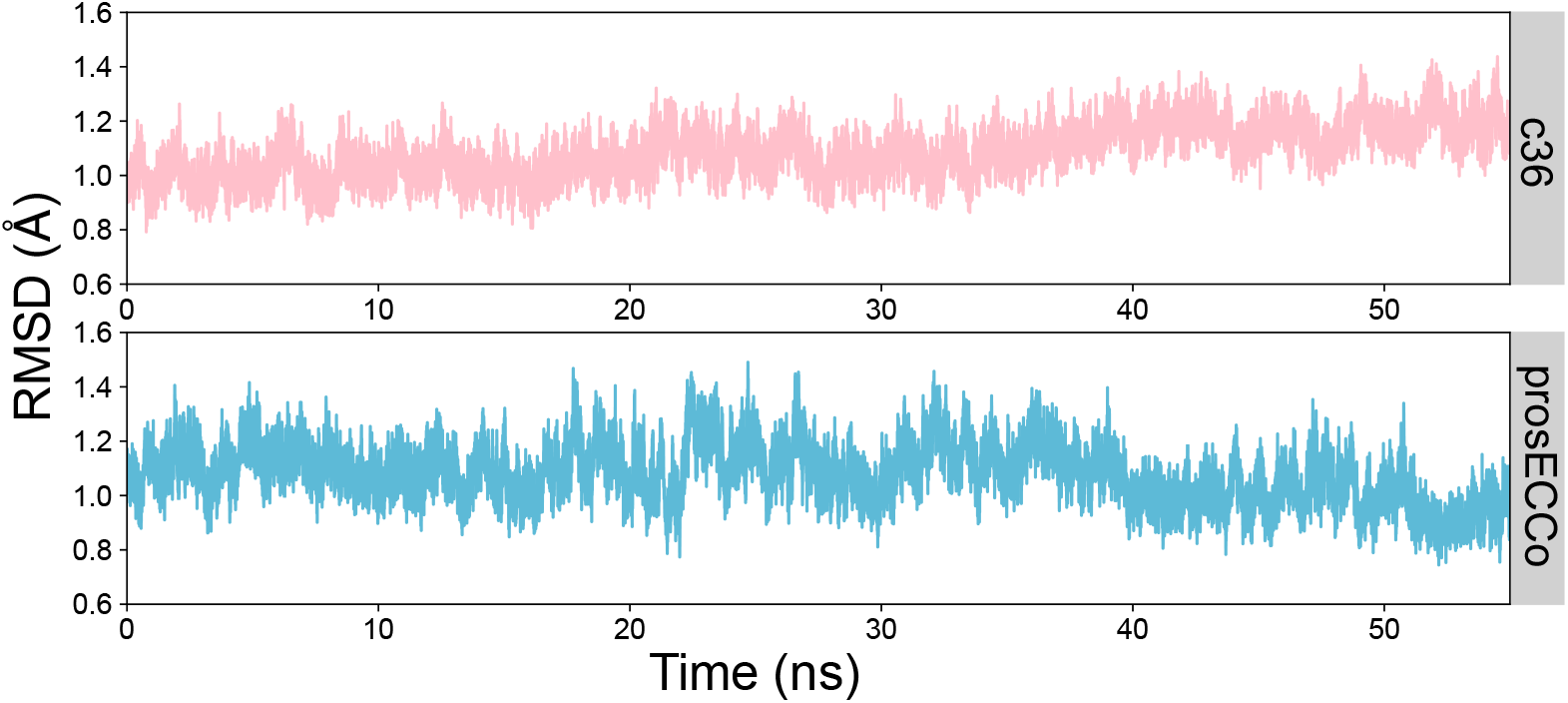
RMSDs of the protein backbone in simulations of the whole protein embedded in a lipid bilayer without backbone position restraints applied. The protein deviates by < 2.0 Å compared to the crystal structure and therefore justifies omitting the use of backbone restraints in simulations of the whole protein.

**Figure S2:**
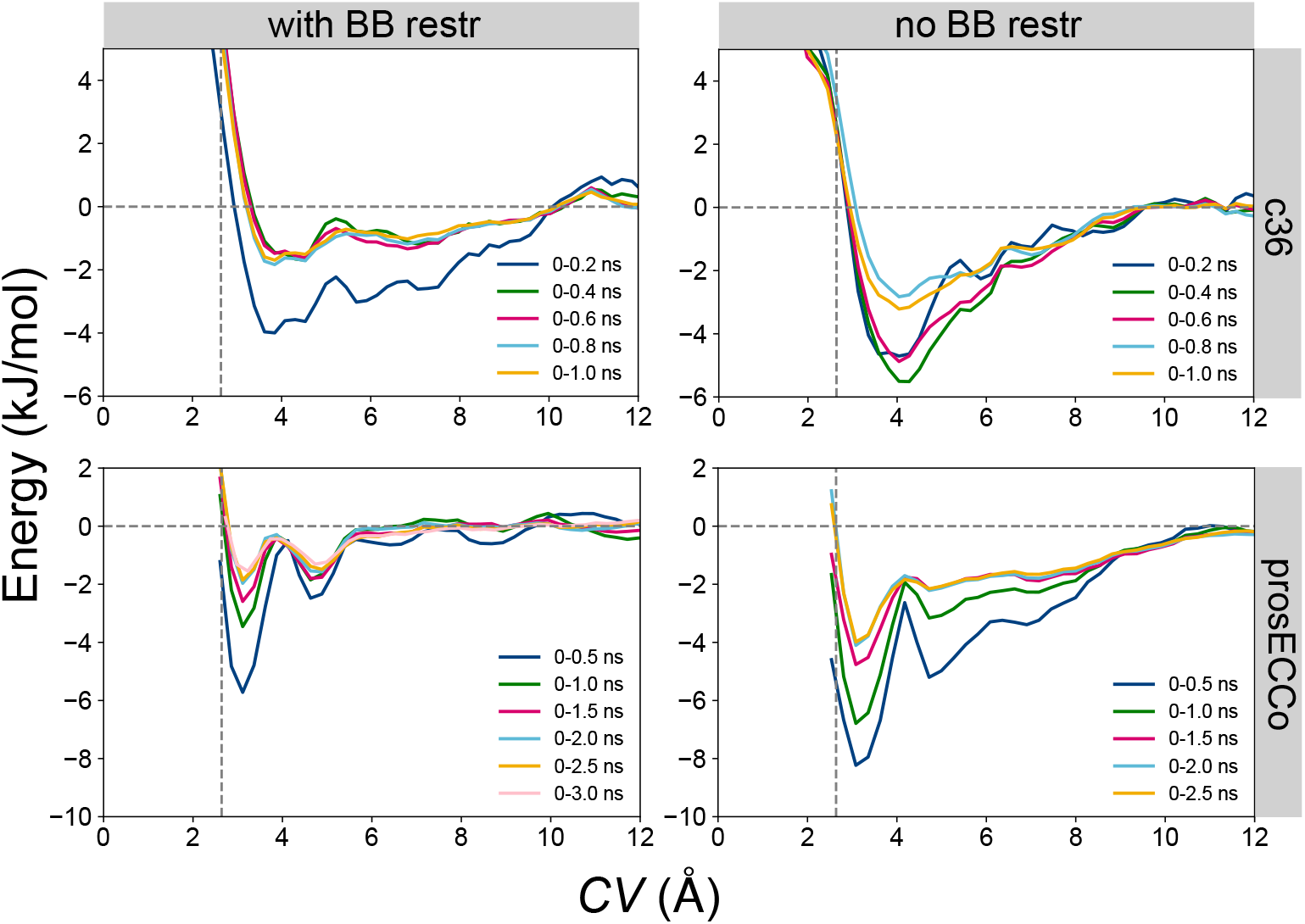
Cl^−^ PMF convergence analysis using the CHARMM36 (c36) and prosECCo force fields with and without the use of protein backbone restraints. Convergence analysis was performed by calculating 0.2 ns and 0.5 ns sampling blocks over the sampling time for of 1.0 ns and 2.5 ns for c36 and prosECCo respectively. The vertical grey dashed line represents the position of the crystal structure binding site in terms of *CV*.

**Figure S3:**
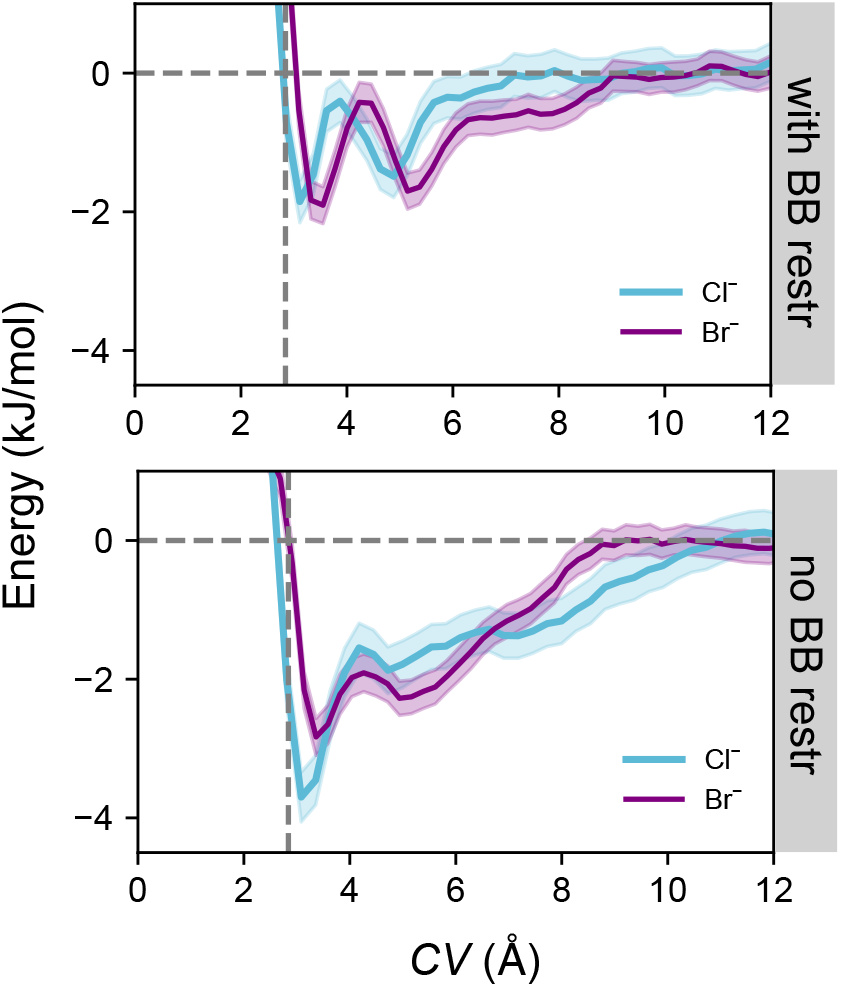
Potential of mean force profiles for a Cl^−^ and Br^−^ ion moving away from the binding site into bulk solution employing the prosECCo force field. Protein backbone restraints were not applied to the simulations. The horizontal grey dashed line represents the free energy of the ion in bulk solution and the vertical grey dashed line is representative of the location of the corresponding ion in the resolved crystal structure.

**Figure S4:**
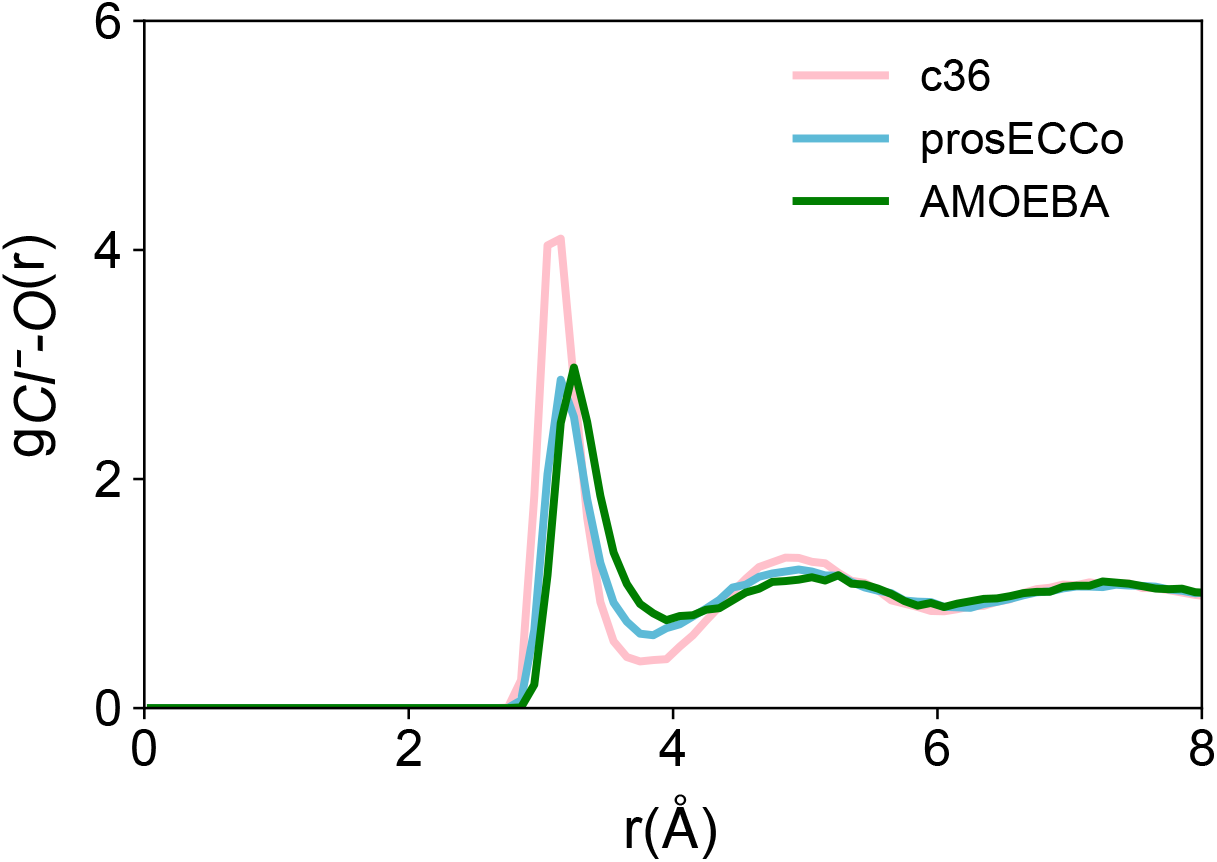
Radial distribution functions (RDFs), *gCl^−^-O* (r), of water oxygen atoms around a Cl^−^ ion in bulk solution with the c36, prosECCo and AMOEBA forcefields.

**Figure S5:**
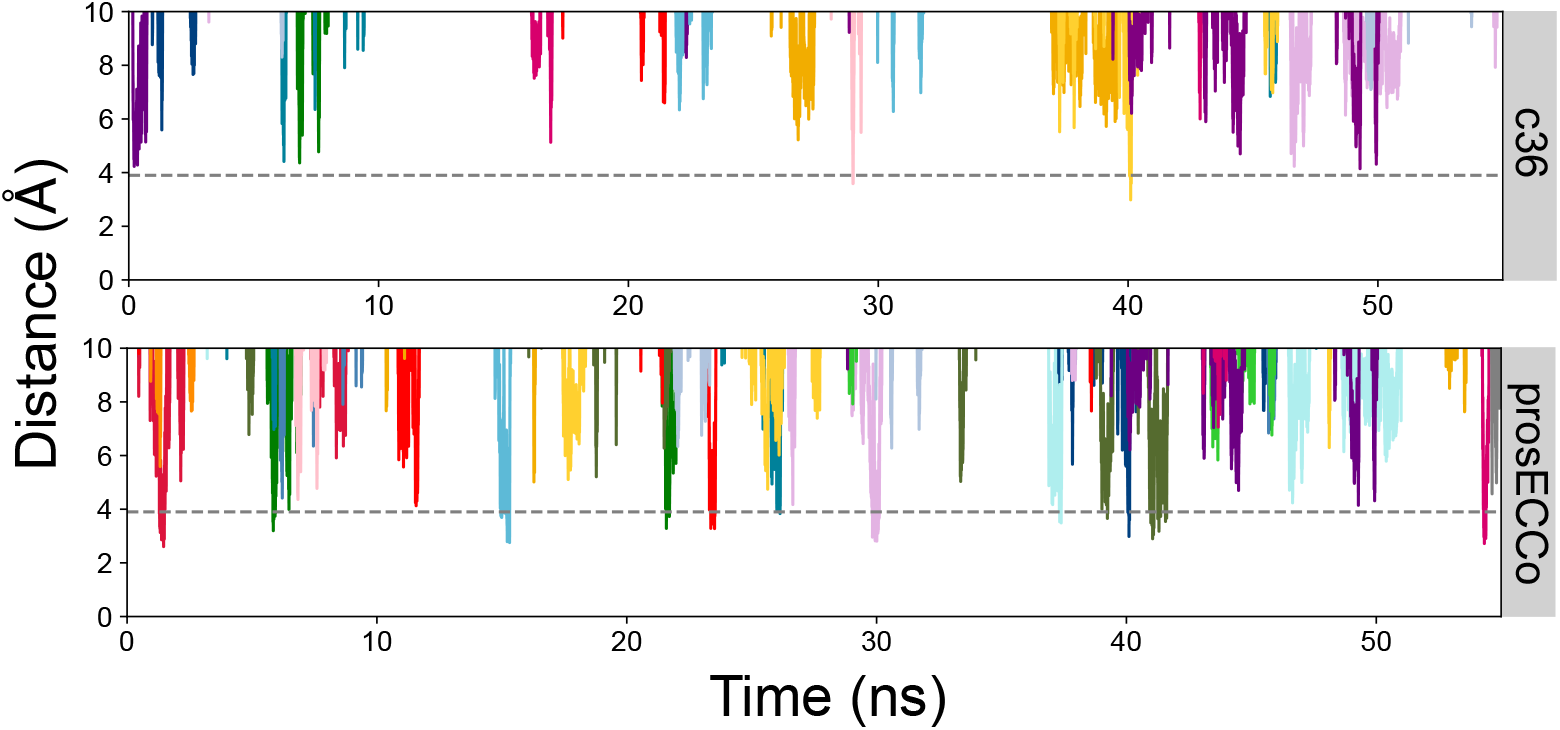
Ion-distance plots between Cl^−^ ions to an aliphatic hydrogen atom of A44 within simulations of the whole protein embedded in a lipid bilayer with the c36 and prosECCo forcefields. Each color represents an individual ion trajectory, plotting only the ions that initially come within a distance of 5 Å to the binding site.

**Figure S6:**
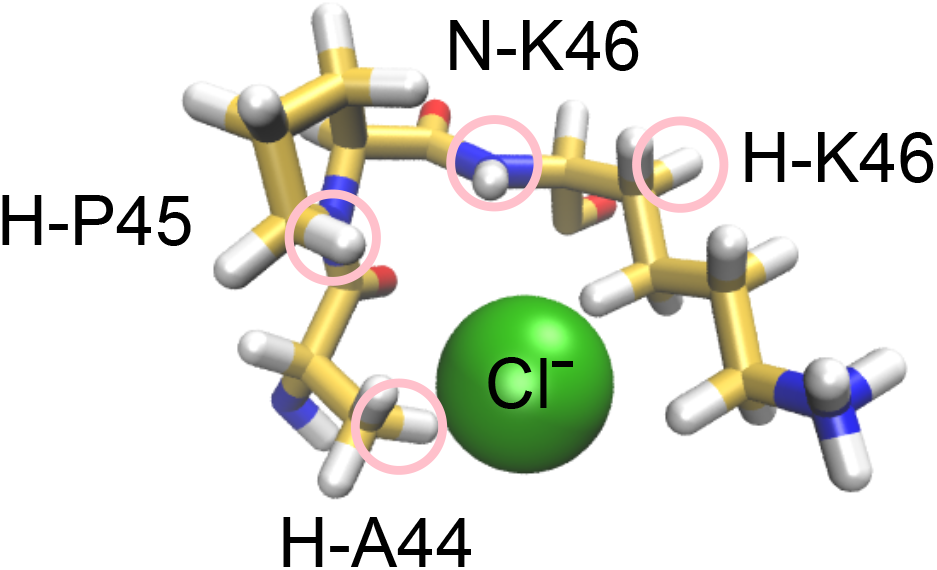
Snapshot of a Cl^−^ ion in the binding site. This is an example of where the K46 tail encloses the ion to form a deeper concave shape that facilitates ion binding.

**Figure S7:**
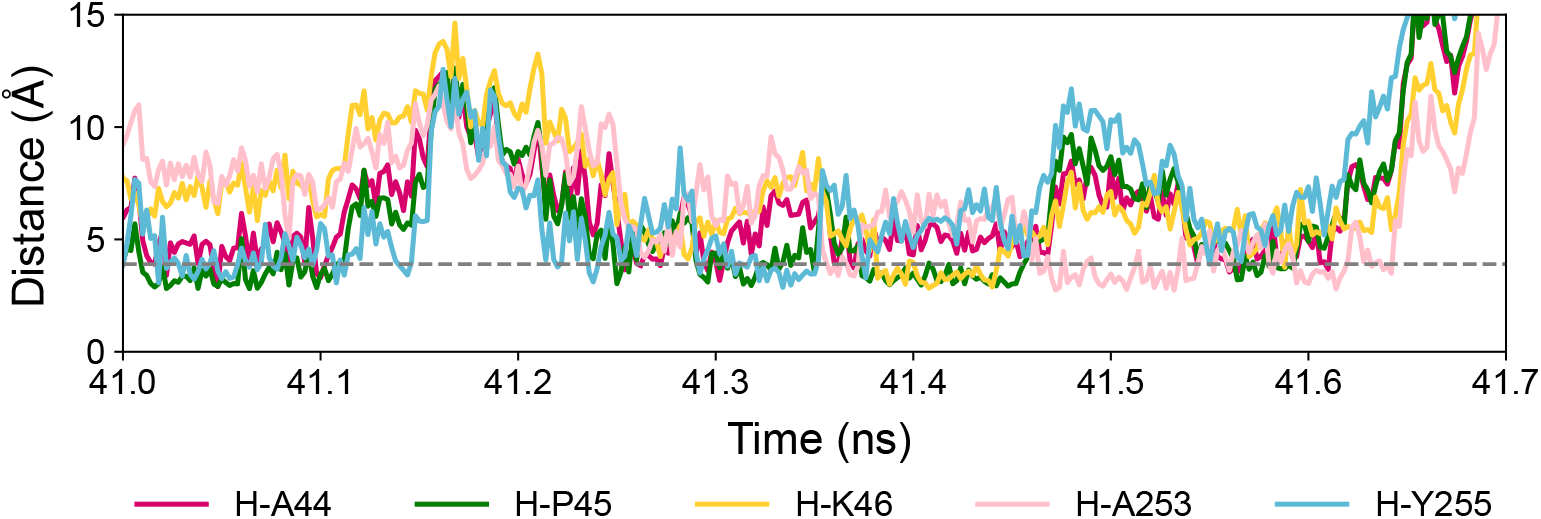
Ion-distance plot as a function of time for a Cl^−^ ion in the binding site and residues from a nearby loop. The ion distances were measured from simulations of the whole protein embedded in a lipid bilayer using the prosECCo force field. The Cl^−^ is bound to the site (contacts H-A44, H-P45 & H-K46) then dissociates (at ~ 41.5 ns) and forms interactions with residues from the nearby loop (H-A253 & H-Y255). Cl^−^ rebinding can be observed (at ~ 41.6 ns) and the ion is then released into bulk solution (at ~ 41.7 ns).

